# MAIA: An open-source, modular, bioreactor for cities

**DOI:** 10.1101/2024.05.03.592196

**Authors:** Andres Rico, David Kong, Kent Larson

**Affiliations:** MIT Media Lab, Massachusetts Institute of Technology Cambridge, Massachusetts, USA

## Abstract

This work presents the design and fabrication of MAIA, an open-source, modular, low-cost, and portable bioreactor for democratizing the development of synthetic biology based projects for urban settings. The integration of open-source synthetic biology (synbio) tools in a city’s infrastructure planning and design is crucial for addressing the great challenges related to urbanization. Synbio tools have great potential to help us complement our current sensing and actuating urban infrastructure. The MAIA reactor controls bacterial growth variables, making it suitable for cell-based experiments while reducing the need for expensive laboratory equipment. Its low-cost and open-source design allow for easy replication and modification, making it accessible to a broader audience. Its portability makes it suitable for use outside of traditional laboratory settings. We qualitatively and quantitatively validated the reactor’s capability to support cell growth, stimulate gene expression, and act as a creative tool for students and users.

## Introduction

The ways in which we plan, design, build and operate cities have lasting impacts on their environmental and social performance [1–3]. In fact, adequate design and operation of cities can have meaningful impact towards allowing us to address several of the United Nations’ Sustainable Development Goals [4, 5]. In the context of cities, sensing and machine intelligence are tools that have become pivotal in helping us to address urban development challenges by allowing researchers, governments and citizens to discover insights that can be used to improve upon current urban decision making processes [6, 7].

As standard city sensors are now well established and helping us understand diverse aspects about how we use our homes [8–10], offices [11] and public spaces [12] amongst other areas; research is now suggesting that the application of biology-based sensors can complement current city sensing capabilities in a meaningful manner (See Figure 1) [13–17]. Advances in city bio-sensing have shown promising alternatives to understanding new ways in which we can instrument our daily lives and infrastructure [18]. Through cell-free systems, site sampling and live cell biosensing, synbio-based approaches have the potential to help us better understand citizens’ needs, environmental degradation, pollution, and behavioral patterns [19–24]. Despite these great advancements, we still lack tools that can help us to better design and deploy these systems within real world contexts.

**Fig 1.**
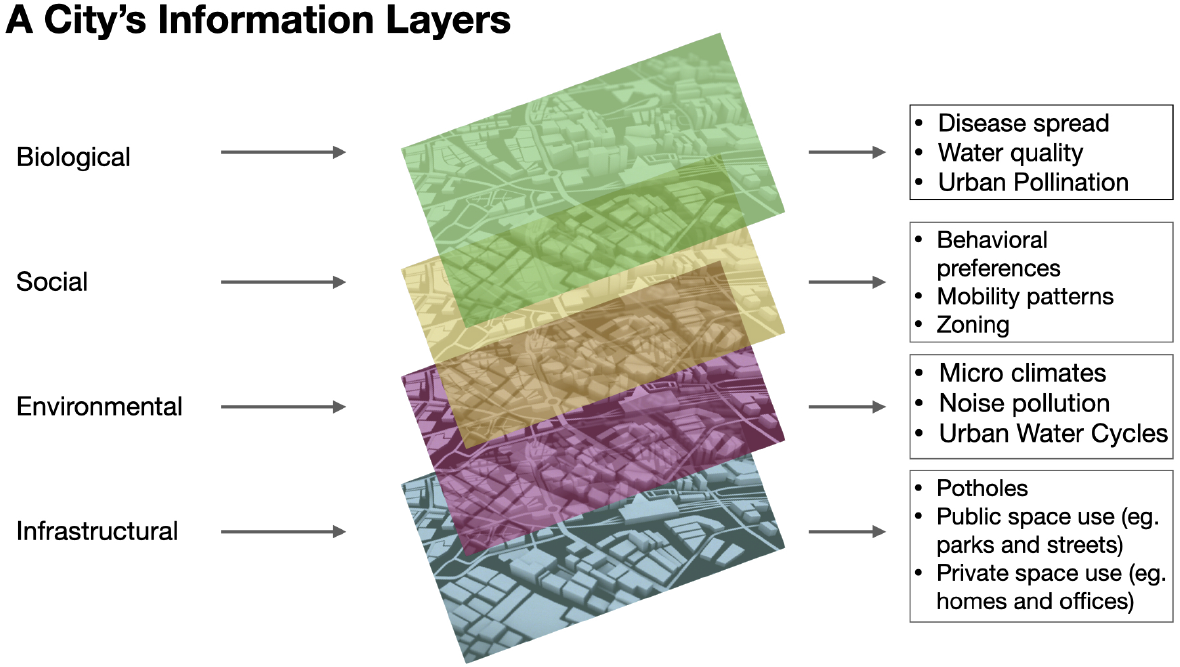
A city’s information layers and examples of insights they provide.

Open-source technologies are playing a pivotal role in democratizing and advancing many scientific fields. Communities and platforms such as Arduino, the Scratch programming language and the Global Fab Lab Network have made great strides to bring the power of computational thinking, robotics and digital fabrication to the most remote corners of the planet [25–27]. Open-source tools allow society to crowdsource problems and solutions that would have taken much longer to be found through traditional top-down scientific processes [28].

The synbio community is not falling behind and is embracing the power of open source to scale access to the field’s development and benefits [29–31]. Recent years have seen the creation of foundations, institutions, devices, and programs that seek to democratize the field. Examples include the BioBuilder Educational Foundation [32], the International Genetically Engineered Machine Competition (iGEM) [33], the Community-Bio-summit, the global How To Grow (Almost) Anything course (HTGAA) [31], and the BioBits Educational Kit [34], amongst others. Moreover, projects like miniPCR [35], AminoLabs [36], Opentrons [37], Foldscope Instruments [38], and Bioexplorer [36] are examples of DIY and educational kits that allow students to engage with essential synbio techniques such as DNA modification, expression, translation and assembly, thermal reactions, and electrophoresis.

More specifically, low-cost reactors and synbio equipment have been of great interest to the scientific community [39–41]. Research has been carried out to develop a large range of low-cost and specialized bioreactors. Some examples include human incubators [42], bioreactors for carrying out field studies in remote or underserved communities [43], laser-cut bioincubators for optogenetic bacterial culture [44], 3D printed microfluidic valves, DNA assemblers and open-source design heuristics [30, 45, 46], flexible cell incubators for affordable, safe, and sterile, hands-on experimentation with live microorganisms [47], microfluidic bioreactors for live-organoid imaging [48] and hypoxia chambers for cell culture [49]. While these reactor designs are breaking barriers in significant ways, there are no open-source and multipurpose devices that can enable researchers, communities and students to run multiple biosensing-related experiments in a portable and low-cost way. Specifically, experiments for developing and deploying novel sensing and information mechanisms for cities and communities.

We demonstrate the development and testing of the open-source MAIA reactor. The reactor controls bacterial growth variables. This is conducive to rapidly deploying cell-based experiments and designs while reducing requirements of expensive laboratory equipment, such as culture incubators and cell stimulation devices. An open-source and modular design allows the device to be easily replicated and modified according to different use cases, needs and contexts. This adds flexibility for it to be easily adapted to fit use cases for city sensing that are limited by user’s experimental needs. For example, users can change the types of stimulation that is given to cells according to different interests (e.g. light stimulation, oxygen stimulation, sound stimulation).

Making the device portable is crucial for taking these systems outside of traditional laboratory settings and democratizing their use. Portability is introduced through a design that is easy to install, self-contained, and lightweight. This unique sum of characteristics results in a device that can be used to further empower the growing space of open-source synbio communities and as a first stepping stone to enabling living cell sensing infrastructure around cities. We evaluate the device by testing (1) its capability to support the growth of cell cultures, (2) stimulate specific gene expression on e-coli cell cultures and (3) evaluating its replicability and usability by student communities. The demonstration of the fundamental characteristics of MAIA enables future researchers to develop applications ranging from sensing biomarkers important for urban health to distributed low-cost scientific exploration tools for community empowerment.

### 1 Bioreactor’s Modular System Design

Bioreactors maintain the stability of specific variables needed for cellular growth [50]. Unfortunately, most reactors act as unified systems, tending to lack flexibility, in the ways in which variables like temperature, vibrations, growth medium, and ventilation are controlled [51]. MAIA modularizes each one of a classic reactor’s functions.

Modularization allows each function or control variable like temperature, ventilation, and growth mediums to be independently assembled and manipulated. Added flexibility lets researchers and students design and attach different stimulation modules according to cell type and experimental design.

Modules are designed in a “LEGO-like” manner to be stacked and ordered according to the type of culture, stimulation, and experiment layout. The design allows for every module to share power and communication lines. Shared power and communication let users run synchronized protocols from the control unit, as shown in Figure 2. The control unit runs the main incubation protocols and communicates with each peripheral module through the I2C protocol, giving it the capability to maintain control of the multiple, in-sync modules that the reactor has the potential to have.

**Fig 2.**
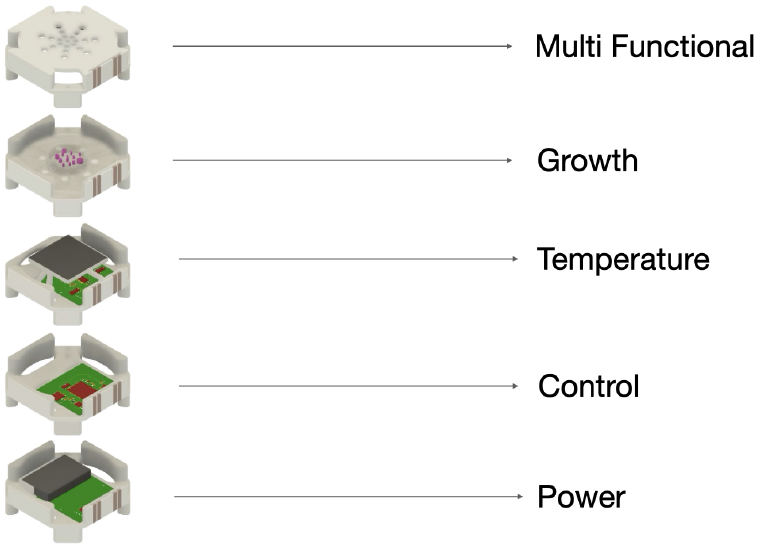
Modular, portable and stackable architecture.

The mechanical design of the modules is standardized so that it can be easily manufactured and replicated by using commonly available fabrication techniques and components. Fabrication of each component is carried out strictly with tools found in the Fab Lab Foundation’s component, material, and machine inventory to keep distributed replication viable. Additionally, the designs are simplified and have physical “poka-yokes” that allow modules to functionally attach without the need for careful adjustment or alignment. It is relevant to note that the complete set of 3D part files, bills of materials, pcb designs, embedded programs, and instructions used to build and program a MAIA reactor can be accessed within this public repository [52].

#### 1.1 Base Structure Module

The base module is the main building block for the system. It acts as the foundation for creating specialized modules. Every module described below uses the base structure and adds different accessories, circuits and components to modify its function. This is the module on which the open-source community would add new functions and modify experimental set ups. The module was designed so that it can be manufactured by 3D printing it on commercially available 3D printers. The current implementation used the Prusa i3 MK3S+ [53] and Formalsb’s Form3 [54] printers to test assembly tolerances and material properties for black and white PLA, and for Formlab’s proprietary Clear Resin V4 [55].

The design can hold two printed circuit boards (PCBs), one on the top side and one on the bottom side. It also has four philipps-like cross plugs (inlets) that can be used to attach accessories that enhance function for the base design such as peltier trays for thermal modules, PCB fasteners, micro-pumps, ventilation lids, etc. Figure 3 shows the manufactured module on three different materials, white PLA, black PLA and clear resin, along with some of the proposed accessories, growth mediums, cell stimulators and PCB’s mounted on it. Noticeably, the same design can be used for a wide range of structural and functional purposes within the reactor’s construction.

**Fig 3.**
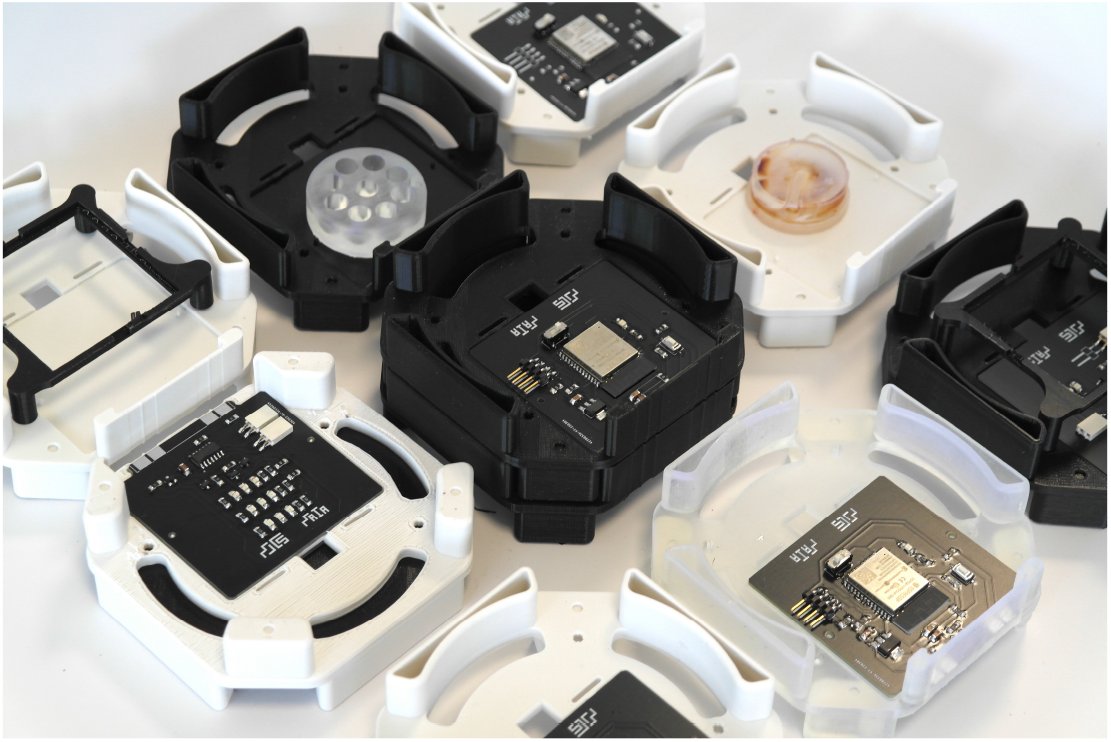
Base modules printed on different materials with PCB’s, growth medium and accessories mounted on them. The image illustrates the flexibility of the module and its capability to be for the different functions that the reactor is required to perform.

#### 1.2 Power Module

The power module supplies power to all of the stacked modules. The design has a standard JST female battery plug. Battery capacity can be varied according to experimental designs. The board also allows for connecting a USB cable’s leads connected to a standard 5V power supply such as the ones found on commercial computers and widespread phone chargers. The power supply can be changed depending on the specific reactor need or experiment being run at any given time. Modifications can be made to the module if the experimental setup contains more modules and thus requires larger amounts of power. In this case, higher current supplies can be introduced into the same form factor. All modules are connected to common power and communication lines in a parallel configuration. Figure 4 shows the power module’s PCB design.

**Fig 4.**
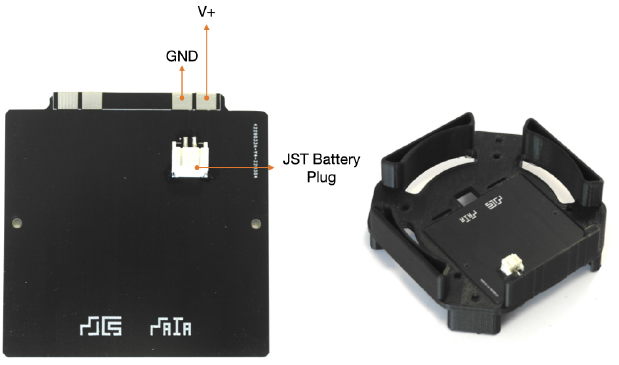
Power PCB module.

#### 1.3 Mother Module

The mother module is the central brain of the reactor. The board is based on the ESP32 microcontroller. The device is WiFi and Bluetooth enabled so that protocols can be programmed externally and commands sent wirelessly to the board. We can program protocols that run on a server and communicate wirelessly with the reactor. This allows the device to be used for a broad range of lab experiments that require variable adjustment over long periods of time.

The motherboard carries out communication with all other modules to check sensor and actuation status of all peripheral boards. The module checks the status of all sensors and actuators and ensures that they are aligned with user defined protocols.

Communication with all other boards is carried out through the I2C protocol in which peripheral modules act as responder devices for the motherboard. The board’s communication capabilities can allow MAIA to be integrated into user friendly interfaces for control. Figure 5 shows the module assembled and mounted on top of the power module.

**Fig 5.**
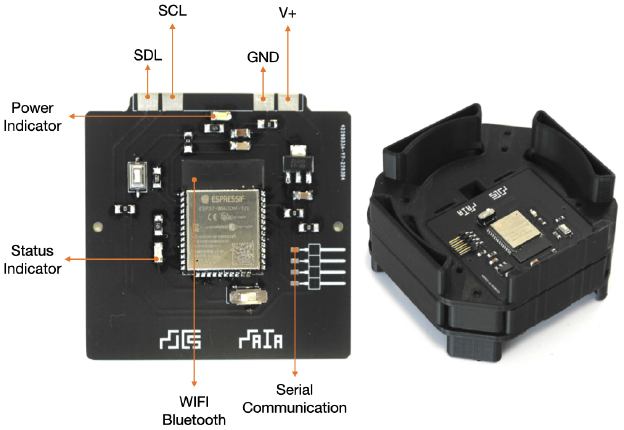
Mother PCB module with WIFI and Bluetooth capabilities.

#### 1.4 Temperature Module

The temperature module is based on an ATTINY 1614 microcontroller. The module controls a peltier thermal element that is in close contact with the growth medium to provide the right temperature for cell reproduction. The peltier effect appears on thermocouples when we pass current through them. One side of the thermocouple looses heat while the other gains it. This allows us to have a reversible and compact heating source for the reactor. The device’s temperature range can be adjusted through the use of 3D printing materials with better thermal characteristics than the current materials. Current materials allow for operation at a maximum of 150° C.

The peltier device is driven by a set of transistors, and a temperature sensor monitors culture temperatures to ensure they are at their set temperatures. The temperature module situates the heating and cooling element (peltier) on a tray that is fixed on the base module. The tray allows for direct transference of heat onto the module that is placed directly above the temperature module. Figure 6 shows the PCB design and peltier tray assembly for the temperature module as it would be stacked on top of the mother module. the assembly tray design can be modified depending on peltier supplier and capacity.

**Fig 6.**
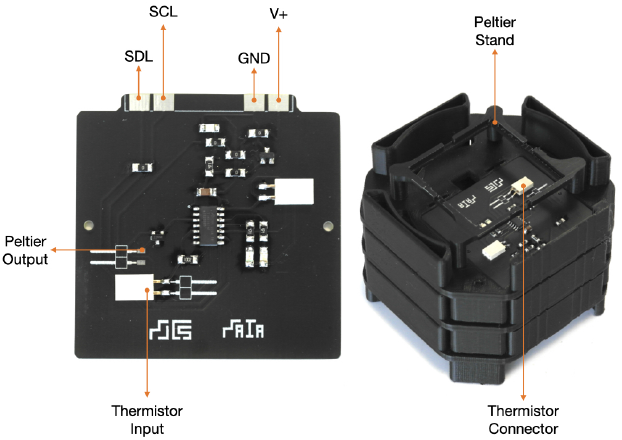
Temperature PCB module.

#### 1.5 Growth Module

The growth module is an empty base module where bacterial cultures and gorwth mediums like agar plates are physically placed. The growth module can house different types of cultures in a cavity that can hold petri dishes with a maximum of 80*mm* of diameter and 14*mm* of height. The current version is placed above of the temperature module so that the Peltier element is in direct contact with the surface of the growth module. This allows bacterial cultures to have an isolated and controlled thermal chamber.

We can control variables like temperature, ventilation, light incidence and heat through the addition of accessories or modules placed on top or below of the growth module. All of this is controlled depending on the desired experimental setup. As illustrated on Figure 7, future versions of the device could have active growth modules that use micropumps and microfluidics to maintain cultures alive for longer periods of time. Complex growth modules could be successively stacked depending on specific experimental requirements.

**Fig 7.**
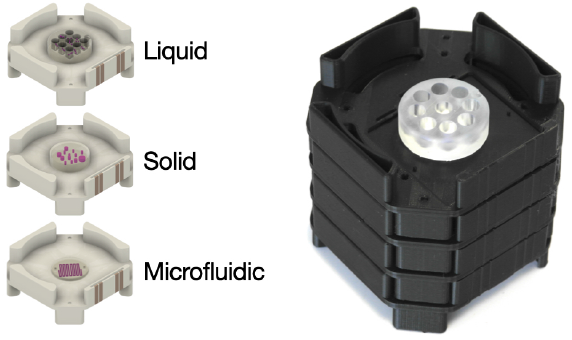
Growth module assembly.

#### 1.6 Stimulation Module

The stimulation module contains the main actuator that will stimulate the cell culture to induce specific gene expressions. Its important to note that the stimulation modules can be specifically built and design according to the cell stimulation that you want to test. One can imagine having light modules, ventilation modules, heavy metal exposure modules, amongst others. For the current validation and demonstration, we designed a light module to test different protein expression circuits that are dependant on the incidence of specific wavelengths on the cells.

As Figure 8 shows, the board is based on an ATTINY 1614 microcontroller which controls three different LED arrays. The arrays shine light in visible wavelengths. The version has capabilities to shine red, blue and green light. The board is mounted on the lower PCB mount of the base module design so that its light shines directly on the bacterial culture. This type of design could be further improved to include other types of light such as infra red or ultraviolet. It could also be to create imaging systems that can monitor cell growth over time through an embedded camera. Figure 8 also allows us to see the scale of the complete reactor assembly and components when compared to a human hand to demonstrate portability.

**Fig 8.**
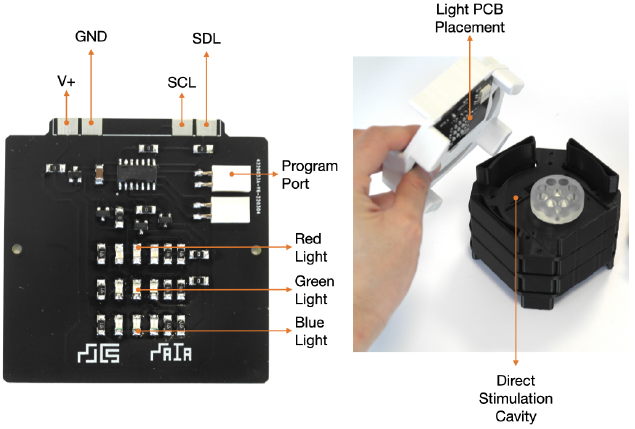
Light PCB module.

**Fig 9.**
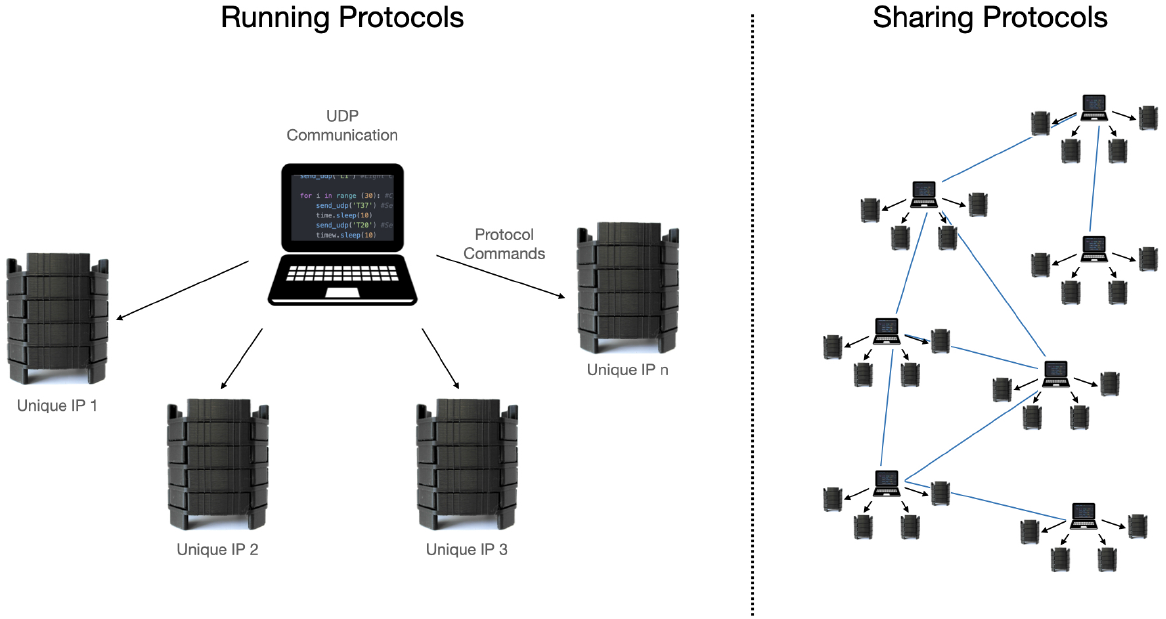
Running protocols on multiple MAIA reactors and sharing protocols to computer and MAIA community networks.

**Fig 10.**
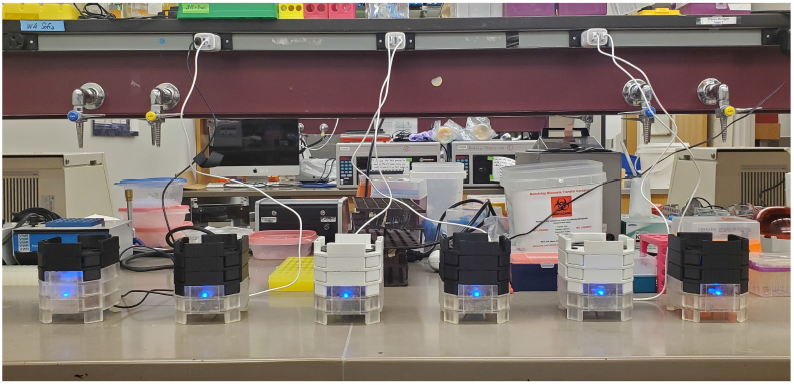
MAIA reactors setup in lab for blue light stimulation experiments.

**Fig 11.**
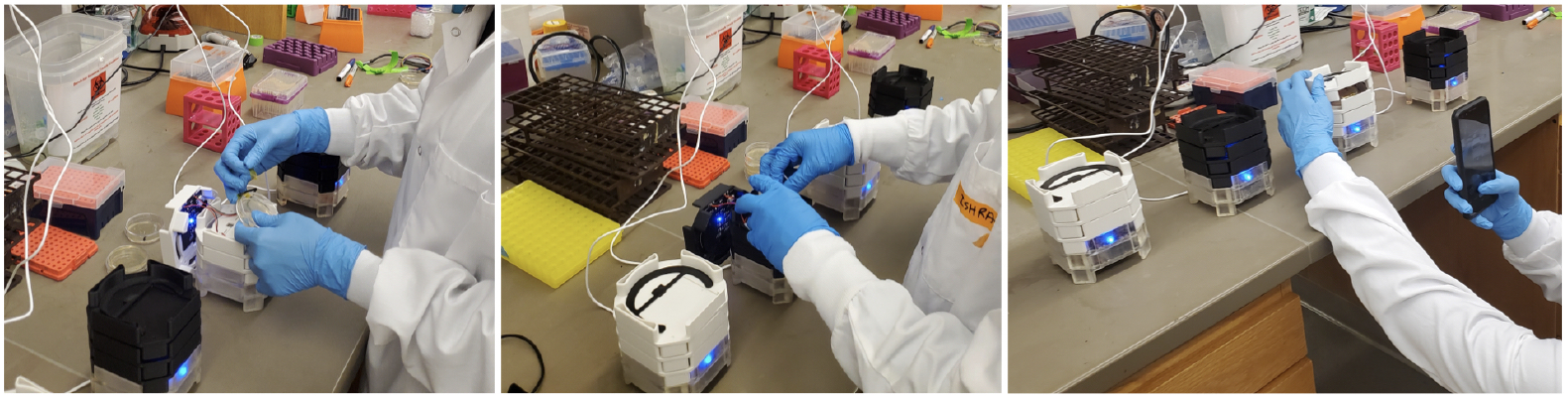
Students using the reactor to set up different experiments.

**Fig 12.**
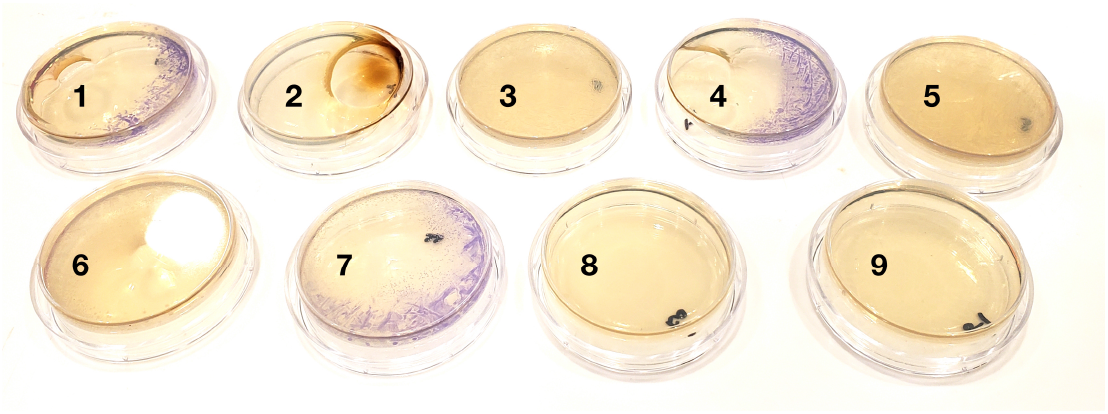
Experiment 1 results for growth and gene expression.

**Fig 13.**
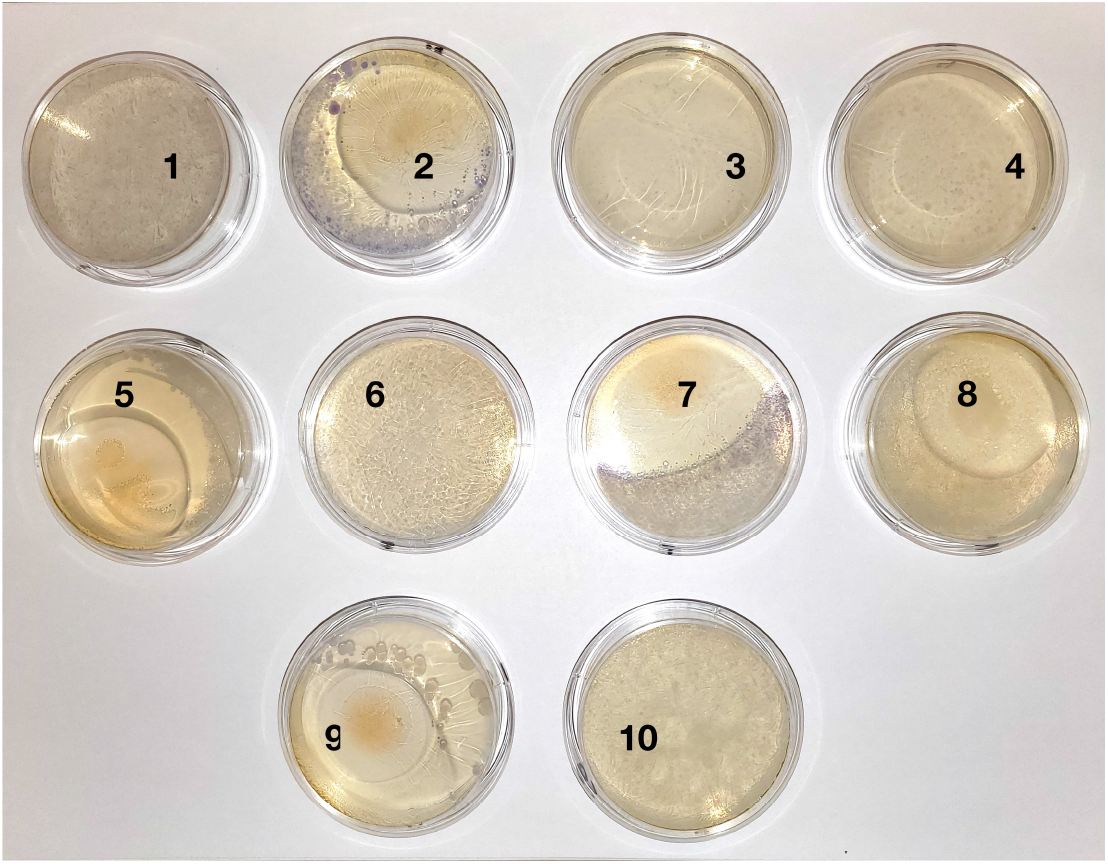
Experiment 2 results.

## 2 Community-shared Bioreactor Software and Protocols

As outlined above, the motherboard is based on a WiFi and Bluetooth-enabled microcontroller that allows users to design and execute reactor protocols directly from a web application. Wireless connectivity allows people with low computational skills to operate the device because no embedded programming is required. this means that higher level languages, such as Python, can be used to operate the device. This is crucial for further lowering barriers of entry to using the device.

In addition, peripheral modules are designed to only send their current sensor and actuation states to the motherboard, and the motherboard will return commands for changing those states according to the user-specified protocol. The device employs user datagram protocols (UDP) to communicate with the motherboard’s WiFi chip. The chip sets a fixed IP address when it connects to the internet. The address is used on a remote server to send protocol commands to it.

Commands can be set up within python scripts. The scripts can then be combined to produce community tested protocols that can be shared and executed on multiple reactors. for example, a student could create a protocol that oscillates the temperature in the reactor for a specific experiment and share it for it to be to other use cases that need temperature oscillation. This is a key characteristic that allows the device to take advantage of open-source dynamics for community protocol generation and testing.

Please reference supplementary material for information on how to construct commands for operating the reactor.

### 2.0.1 Module Protocols

#### Temperature Module Program

The temperature board uses a closed-loop control sequence that allows it to maintain a desired temperature. The program constantly reads the thermistor’s value and activates the Peltier element as needed. The Peltier element is activated to cool down or heat up until a desired temperature value is reached. The last desired temperature that the device receives is stored on internal memory so that any momentary power outage does not disrupt a protocol.

#### Light Module Program

The light board only has actuation elements on it. Its three outputs switch the light arrays on and off. The board constantly listens to updates on light commands, stores new values and executes them by switching specific LED’s on and off.

## 3 Experimental Design

The reactor’s validation is done through two experiments. The first experiment uses multiple MAIA bioreactors to evaluate growth and stimulation of a bacterial blue light sensing system. The second experiment is done within a hands on workshop. The workshop helps to evaluate performance of the device when operated and assembled by students and the MAIA bioreactor’s potential for being used for a wide range of applications.

### 3.1 Light Responsive Biosensor Design and Construction

We evaluate the performance of the device by growing a solid agar based e-coli culture. To do so, we run two experiments. One of the experiments was done in preparation of the workshop and the second one was carried out during the workshop with student participation.

The experiments validate bacterial growth and gene expression correlated with specific stimulation conditions. We place the bacterial culture on agar plates and grow them within different reactors. Each of the reactors uses different settings for temperature and light stimulation.

The experiments are run with an e-coli K12 strain transformed with a blue light sensitive genetic circuit described in [56]. The plasmid used contains a circuit with a photoreceptor and a chromoprotein reporter. Blue light activates the photreceptor, activation leads to production of blue pigmented proteins. The plasmid uses a combination of the mUAV plasmid [57], which contains the AmilCP gene, and the pDawn plasmid [58], which contains the YF1/FixJ light system. According to [56], the YF1/FixJ is first inserted into a pET-21(+) vector via Twist Bioscience. Restriction cloning is then used to insert the amilCP gene into the new plasmid.

Cell cultures were first grown as liquid cultures in a commercial incubator. The cultures were then transferred onto 30*mm* diameter Petri dishes with Ampicillin and a standard LB agar mix from Fisher Scientific [59] for running all of the experimental setups. It is relevant to note, that the 18*mm* tall dishes were used due to supply chain issues but the reactor’s design is made for 15*mm* tall dishes.

### 3.2 Student Workshop

The device was tested in the context of a hybrid workshop imparted for the How To Grow (Almost) Anything class [31]. The workshop was held at the MIT Campus in Cambridge, MA, USA. The goals of the workshop include (1) introducing students to the MAIA reactor as a platform, teaching them how to build and use it and (2) evaluating performance characteristics for the device when operated by the students. The workshop had eight in-person student participants and eight online participants. All in-person students were able to directly interact with setting up reactors and experiments. Figure 14 shows images of students engaging with the bioreactor during the workshop.

**Fig 14.**
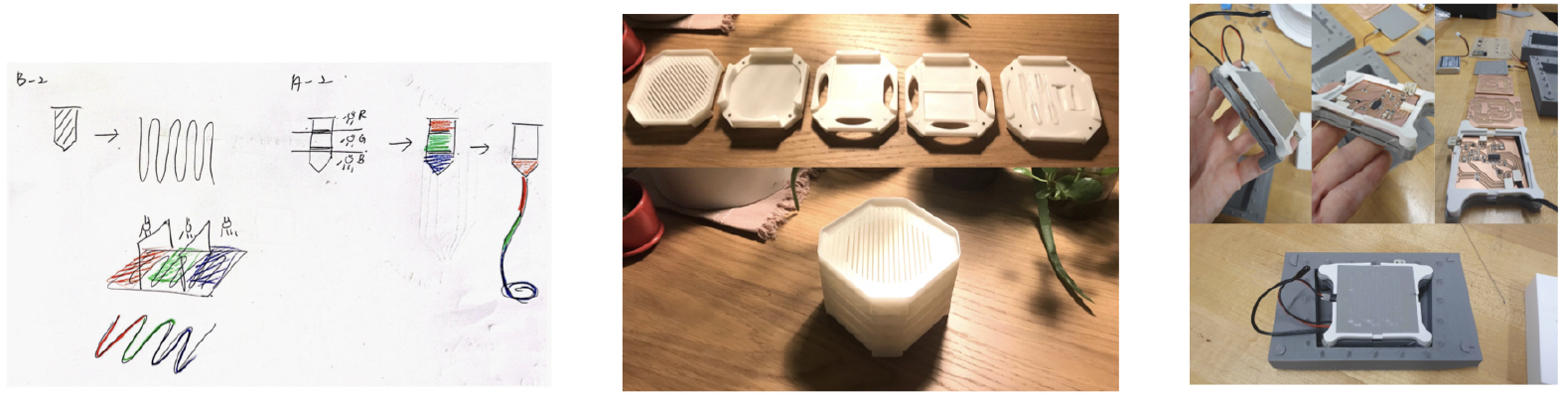
Student sketches and prototypes.

The workshop had three sections. The first section of the workshop introduced basic concepts of bioreactors, and specific information about how MAIA is built, how to replicate it and how to use each one of its components. This section also included discussion on the bio-ethics and possible policy changes that need to be addressed for keeping the deployment of bioreactors around cities safe. The second section corresponds to the use case implementation with light sensitive bacteria. Students define experimental setup according to the bioreactor’s capabilities and collectively indicate a hypothesis for each one of the tests. Lastly, the third section was a hands-on experience setting up six MAIA reactors and ten experiments to run. Six of the experiments were setup within MAIA reactors and the four others were set up as positive and negative controls within commercial incubators. Students defined variables, set them on each reactor, prepared their samples and installed the agar-filled Petri dishes inside of the units.

## 4 Results

### 4.1 Cell Growth and Gene Expression

The first experiment used seven reactors to evaluate cell growth and performance of the light and temperature modules. Table 1 shows the experimental setup used on each reactor. The bacterial culture was incubated using the e-coli strain described above. 25*µL* of culture were put into each of the agar plates and left within the reactors for 24 hours.

**Table 1.**
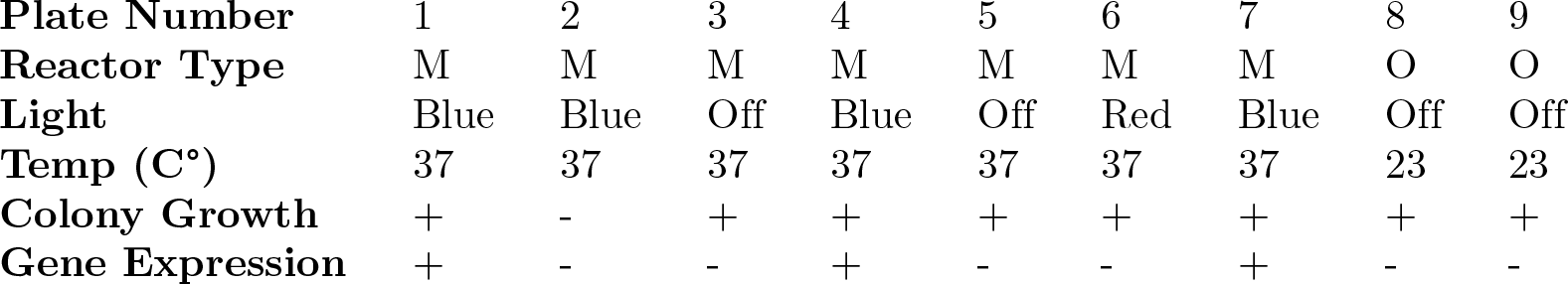
Experimental setup and results for the first experiment.

Results showed that eight out of the nine experiments were successful. Reference experiment table 1 and Figure 13 to see results. The cell cultures reacted to the growth and stimulation devices as expected. Only one of the reactors malfunctioned. The cause is though to be a problem in the temperature module coming from a faulty temperature sensor connection.

The experiment evaluates cell ground and gene expression on bioreactor. Reactor key: M = MAIA reactor, O = Outside (no incubator), I = Commercial incubator.

The experiment’s results allow us to conclude that the reactor is able to grow cell cultures and vary the light stimulation that it can give to evaluate the performance of a light sensitive system on e-coli ^1^. Cell cultures grew as expected and gene expression was seen on all of the experiments that had blue light stimulation. Experiments with stimulation of other wavelengths correctly showed cell growth but no gene expression could be seen.

### 4.2 Student Use of Reactor

The second experiment was carried out within the context of the workshop. The bacterial liquid culture was prepared as outlined above. For this experiment we used 10*µL* of culture on each Petri dish and allowed cultures to grow for 48 hours. Students were tasked with designing their own experimental setups and variables. Table 2 outlines the experimental setups chosen by students. This batch of experiments included positive controls grown on the lab’s controlled temperature chambers.

**Table 2.** Experimental setup and results for the second experiment.

|  |  |  |  |  |  |  |  |  |  |  |
| --- | --- | --- | --- | --- | --- | --- | --- | --- | --- | --- |
| **Plate Number** | 1 | 2 | 3 | 4 | 5 | 6 | 7 | 8 | 9 | 10 |
| **Reactor Type** | I | M | M | O | M | O | M | M | M | I |
| **Light** | White | Blue | Blue | Off | Green | White | Blue | Red | Off | Off |
| **Temp (C°)** | 37 | 37 | Room | Room | 37 | Room | 42 | 37 | 37 | 37 |
| **Colony Growth** | + | + | - | - | + | - | + | + | + | + |
| **Gene Expression** | + | + | - | - | - | - | + | - | - | - |

The experiment evaluates cell growth and gene expression on bioreactor carried out by students. Recator key: M = MAIA reactor, O = Outside (no incubator), I = Commercial incubator.

Results show that nine out of the ten experiments worked as hypothesized. We can see that bacterial growth is less dense than than growth from experiment one. This is because we used a smaller concentration of cells for each of the plates, This is also reflected on the intensity of color for the colonies.

Both experimental results allow us to validate that the reactor’s temperature control and growth modules, maintain an ideal temperature to allow for cells to grow. The light isolation and stimulation also had positive results on experiment two as blue protein pigments can be observed on cultures stimulated with blue light.

With this experiment we can also evaluate that the reactor can be successfully operated by students as all of the cultures setup by students showed bacterial growth. In the results, we see variation in uniformity and density of cultures, this is attributed to the variability that is introduced as students set up their own pipettes, load the cells onto the dish, and insert each dish into a reactor. With this we can conclude that both of the light experiments allow us to verify correct performance of the device.

Additionally, after setting up their experiments, students in the workshop were asked to envision how they could modify the reactor to their specific interests and project ideas. Students then produced concept sketches and prototypes applying the bioreactor for bio-production of tinted silk, biocementation and bacteria based textile sensors.

Figure 14 illustrates some of the concepts, sketches and prototypes that were made by students. These are clear examples of the creative benefits that these types of bioreactor designs could unlock for cities at different scales when democratized across disciplines, domains and communities.

## 5 Discussion

Creating technological solutions that can be easily scaled and used by communities that have very contrasting needs, skills and resources is critical within the frame of democratizing technology and keeping its benefits safe and distributed. Specifically when addressing urban challenges, this means that we should shift our focus from only creating smart devices to enabling smart communities. With this shift, deploying technologies becomes less about finding centralized ways to impose technological solutions and more about giving communities skills, tools and safe regulatory frameworks for them to develop their own unique solutions that respond to their specific needs. Critical to achieving this goals, is the development of devices that are able to adapt and be expanded upon the creativity and needs of communities with different disciplines, backgrounds and needs. We view the open-source, modular and portable essence of the MAIA reactor as a clear embodiment of a device with these characteristics.

In the future, we can envision communities that could use these types of bioreactors to actively monitor for undesired metals in their water infrastructure or to collectively monitor the spread of a disease. Similarly, researchers could monitor the patterns in the spread of pollen around urban areas so as to understand how urban design can impact behavior patterns of insect species and their environments. Devices could also be built to measure the impact that different urban micro climates could have on a range of human or non human cells. These types of analysis could ultimately lead us to understand ways in which we can purposefully design cities to have a more balanced relationship with the natural environment.

In a broader view, within urban settings, these types of devices could also allow for decentralized bio-production of compounds that could be used to combat health threats or improve hyper-localized food production processes. This could lead to significant changes in how we produce and consume resources in urban environments. Furthermore, our improved understanding of biological processes within cities could serve as inspiration for the implementation of dynamic zoning algorithms, decentralized mobility coordination systems, and self-regulating heating and cooling infrastructure for buildings. Exploring and utilizing the biological layers of cities has significant potential to help us learn not just how to improved city design, but also more effective ways of living and managing them.

As these types of devices start to become more ubiquitous around cities, it is critical to note the importance of proper regulation for them. Specifically regulations that address cross-contamination or malicious system design. Biosafety regulations already in place for heavy laboratory infrastructure can be adequately adapted for the upcoming waves of open-source, DIY tools and infrastructure. The open-source nature of the device can also be used as an inherent mechanisms of community supervision. As has happened with many digital technologies, community vetting of designs and applications can allow for decentralized control and alignment of these powerful technologies.

In conclusion, the present work outlines the design and validation of MAIA, a low-cost, modular and portable bioreactor. The bioreactor’s design proves to be a unique and valuable addition to the space of low-cost bioreactors as well as for foundations, competitions and devices that aim to democratize Synthetic Biology. The system is evaluated for its capability to grow cell cultures and stimulate them. We also demonstrate how the system can be used by student communities to fit with personal project ideas and research directions. We finally discuss how this growing ecosystem of low-cost, open-source and community oriented devices allows us to envision a future where communities are able to respond to challenges with highly sophisticated technologies built collectively in-situ to better address a growing landscape of challenges in cities around the globe.

## Footnotes

1 Plates that had the light stimulation module on top of them, show uneven cell growth. This is attributed to direct contact to heat from the light module. Contact was enabled by using petri dishes that are not of appropiate size to fit in growth module

## Notes

https://github.com/AndresRicoM/MAIA

### Competing Interest Statement

The authors have declared no competing interest.

